# Cytotoxic chemotherapy potentiates the immune response and efficacy of combination CXCR4/PD-1 inhibition in models of pancreatic ductal adenocarcinoma

**DOI:** 10.1101/2023.12.24.573257

**Authors:** Alexander G. Raufi, Ilenia Pellicciotta, Carmine F. Palermo, Steven A. Sastra, Andrew Chen, Emily Alouani, H. Carlo Maurer, Michael May, Alina Iuga, Raul Rabadan, Kenneth P. Olive, Gulam Abbas Manji

**Author notes:** These authors contributed equally (Alexander G. Raufi, Ilenia Pellicciotta). Please address correspondence to Kenneth P. Olive or Gulam A. Manji.

## Abstract

**Purpose:** The CXCL12-CXCR4 chemokine axis plays a significant role in modulating T-cell infiltration into the pancreatic tumor microenvironment. Despite promising preclinical findings, clinical trials combining inhibitors of CXCR4 (AMD3100/BL-8040) and anti-programmed death 1/ligand1 (anti-PD1/PD-L1) have failed to improve outcomes.

**Experimental Design:** We utilized a novel ex vivo autologous patient-derived immune/organoid (PDIO) co-culture system using human peripheral blood mononuclear cells and patient derived tumor organoids, and in vivo the autochthonous LSL-KrasG12D/+; LSL-Trp53R172H/+; Pdx-1-Cre (KPC) pancreatic cancer mouse model to interrogate the effects of either monotherapy or all combinations of gemcitabine, AMD3100, and anit-PD1 on CD8+ T cell activation and survival.

**Results:** We demonstrate that disruption of the CXCL12-CXCR4 axis using AMD3100 leads to increased migration and activation of CD8+ T-cells. In addition, when combined with the cytotoxic chemotherapy gemcitabine, CXCR4 inhibition further potentiated CD8+ T-cell activation. We next tested the combination of gemcitabine, CXCR4 inhibition, and anti-PD1 in the KPC pancreatic cancer mouse model and demonstrate that this combination markedly impacted the tumor immune microenvironment by increasing infiltration of natural killer cells, the ratio of CD8+ to regulatory T-cells, and tumor cell death while decreasing tumor cell proliferation. Moreover, this combination extended survival in KPC mice.

**Conclusions:** These findings suggest that combining gemcitabine with CXCR4 inhibiting agents and anti-PD1 therapy controls tumor growth by reducing immunosuppression and potentiating immune cell activation and therefore may represent a novel approach to treating pancreatic cancer.

## INTRODUCTION

Immune checkpoint blockade (ICB) has dramatically improved outcomes for individuals diagnosed with multiple tumor types, however these agents have largely been ineffective in pancreatic ductal adenocarcinoma (PDAC) [1-9]. It is hypothesized that this is not exclusively due to PDAC-intrinsic resistance to ICB, but also due to a highly immunosuppressive tumor microenvironment (TME), which is partly induced by tumor-resident stromal cells [10-12]. Observations that high T-cell infiltration and the presence of tertiary lymphoid structures correlate with longer survival of PDAC patients suggest capacity for immunogenicity [13, 14]. Furthermore, although PDAC harbors a moderate mutation burden, this has been shown to sufficiently create highly immunogenic neoantigens [15]. Indeed, these hypotheses are supported by clinical data from an early phase trial in which adjuvant autogene cevumeran, a personalized RNA neoantigen vaccine, stimulated expansion of neoantigen-specific T-cells and resulted in improved recurrence-free survival in participants who developed T-cell responses [16]. Contemporary approaches to increasing the efficacy of ICB in PDAC have focused on targeting components of the desmoplastic stroma in an effort to increase T-cell penetration. To this end, targeting signaling pathways directly responsible for T-cell exclusion, such as the CXCL12-CXCR4 axis, is of particular interest [17, 18].

The CXCL12 chemokine, upon binding to the CXCR4 receptor on T-cells, modulates integrin expression, and at high levels inhibits T-cell migration [19, 20]. Results from preclinical studies using the genetically engineered *Kras^LSL.G12D/+^; Tp53^LSL.R172H/+^; Pdx1Cre^tg/+^* (KPC) mouse model recognized promising anti-tumor activity with CXCR4 blockade when used in conjunction with an antibody targeting programmed death-ligand 1 (αPD-L1) [19]. This combination mobilized T-cells into the TME and stabilized KPC mouse tumors in a 6-day study, yielding increased neoplastic cell death in *ex vivo* human PDAC tissue sections [19, 21]. These data led to multiple clinical trials interrogating the role of targeting the CXCL12-CXCR4 axis in combination with ICB as a potential treatment approach to PDAC [22-24].

Early clinical trials evaluating single or combined blockade of CXCR4 and PD-L1/programmed cell death 1 (PD1) in PDAC demonstrated an integrated systemic immune cell activation and local immune cell recruitment [25, 26]. However, these encouraging correlative findings failed to translate to improved patient outcomes in the absence of anti-neoplastic therapy as only 3.4% of participants experienced a radiographic response [22-24]. The addition of 5-Fluorouracil and liposomal irinotecan to BL-8040 (motixafortide; a CXCR4 antagonist) and pembrolizumab (α-PD1) resulted in a modest partial response rate of 13.2% in metastatic PDAC patients who had progressed on first-line therapy [23, 26]. Perhaps this is due to modulation of the TME, induced by first-line chemotherapy, resulting in reduced reliance on the CXCL12-CXCR4 axis for immune evasion. Alternatively, a gemcitabine-containing regimen may be better suited as a chemotherapy backbone. Therefore, a deeper understanding of the underlying biology of the CXC12-CXCR4 axis is needed to advance therapies targeting this pathway.

Using both human and rodent model systems, we report that, in addition to stroma-mediated mechanisms, CXCR4 inhibition modulates tumor immune responses through cell autonomous mechanisms. We developed and utilized an *ex vivo* autologous patient-derived immune/organoid (PDIO) co-culture system to demonstrate that disruption of the CXCR4-CXCL12 axis leads to increased migration and activation of CD8^+^ T-cells. In addition, when combined with chemotherapy, CXCR4 inhibition (CXCR4i) further potentiated CD8^+^ T-cell activation. We explored these effects *in vivo* using the KPC mouse model and found that the combination of gemcitabine, ICB (α-PD1 mAb), and CXCR4i (triple therapy) resulted in increased cell death, decreased cell proliferation, and a more favorable tumor immune microenvironment as evidenced by an increased CD8^+^/regulatory T-cell (Treg) ratio and increased infiltration of natural killer (NK) cells. Furthermore, these treatments altered the composition and spatial relationships of T-cell and tumor cell populations. Finally, the combination of gemcitabine, CXCR4i, and α-PD1 therapy extended survival in the KPC mouse model. Taken together, these data support the use of gemcitabine-based chemotherapy together with the combination of CXCR4 modulating agents and ICB and demonstrate a novel facet to the immunomodulatory effects of this actionable pathway.

## RESULTS

### CXCR4 inhibition enhances effector immune cell activation and migration in autologous patient-derived immune/organoid (PDIO) co-cultures

To interrogate the effects of CXCL12-CXCR4 signaling modulation on malignant epithelial and immune cells, we harvested paired sets of patient-derived tumor organoids (PDTOs) and peripheral blood mononuclear cells (PBMCs) for use in coculture and transwell migration assays. Five patients with PDAC undergoing surgical resection at Columbia University Irving Medical Center/New York Presbyterian Hospital consented to having both their blood and tumors collected (CUIMC IRB AAAR9431). Once PDTO and PBMC lines obtained from these patients were established, we confirmed CXCL12 and CXCR4 expression. PDTOs were found to secrete varying amounts of CXCL12 **(Fig. S1a)** and CXCR4 was expressed on a subset of PBMCs **(Fig. S1b, left)**. Pre-treatment with AMD3100, an FDA approved CXCR4 inhibitor, prevented the binding of a fluorescently labeled αCXCR4 antibody whose epitope competes with the AMD3100 target on CXCR4 **(Fig. S1b, right)**. These findings are consistent with observed CXCR4 and CXCL12 expression patterns in human tumor tissue and PBMCs **(Fig. S1c).**

We next assessed whether the established PDTOs could activate autologous PBMCs. We quantified the percentage of activated T-cells (CD3^+^CD8^+^IFNγ^+^ PBMCs) before and after co-culture with matched PDTOs and found that co-coculturing PBMCs with matched PDTOs resulted in a significant increase in activated T-cells (**Fig. 1a**). Notably, this effect was not seen when PMBCs were co-cultured with tumor-adjacent patient derived organoids (Adjacent), suggesting a possible dependence on neo-antigen presentation (**Fig 1b**). To investigate this further, we pretreated PDTOs with HLA neutralizing antibodies and noted a 2.3-fold reduction in T-cell activation as compared to controls (**Fig 1c**). Conversely, treatment with cytotoxic chemotherapy (gemcitabine) increased T-cell activation by 1.7-fold (**Fig 1d**). Taken together, these results demonstrate that co-culturing PDTOs with autologous PBMCs *ex vivo* results in increased T-cell activation which may be dependent on antigen presentation and further enhanced with gemcitabine.

**Fig. 1.**
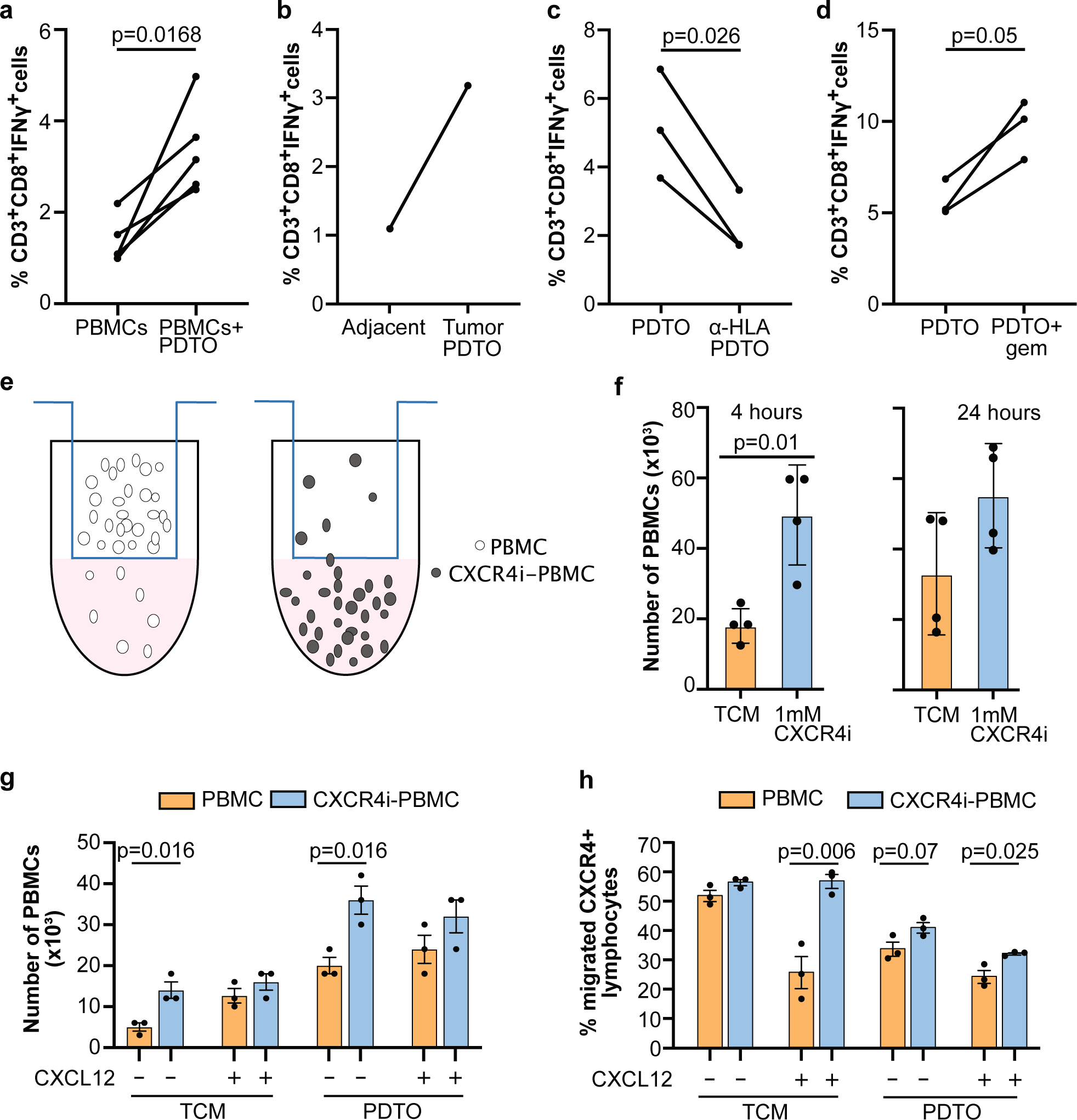
Peripheral blood mononuclear cell (PBMC) migration and activation increases in the presence of PDAC patient derived tumor organoids (PDTOs) and is further enhanced with CXCR4 inhibition. **a**, Induction of T-cell activation, measured by percent of IFNγ expressing CD8^+^ T-cells, in PBMCs alone or after co-culture with autologous PDTOs for 14 days from five distinct PDAC patients. Data are presented as an average of three (n=3, patients hT56 and T19) independent experiments, except for CUMC3, CUMC4, and T18 (n=1; each). Each line corresponds to one patient. **b,** T cell activation after 14 days of co-culture with PDTOs or patient derived tumor-adjacent organoids (Adjacent) from patient CUMC4 (n=1). **c,** T cell activation after 14 days of co-culture with PDTOs from patient T19 (n=2) and T18 (n=1) in the absence or presence of HLA-ABC and HLA-DR/DP/DQ blocking monoclonal antibodies. Each line corresponds to one experiment **d,** T cell activation after 14 days of co-culture with PDTOs from patient T19 (n=2) and T18 (n=1) pretreated for 24 hours with either vehicle or 10 mM gemcitabine. **e,** Schematic representation of the autologous PBMC migration model system using the Boyden transwell assay. **f,** Of the 50,000 PBMCs placed on the top chamber, quantitation of the migrated PBMCs from patients T12 and WT4 treated with PBS in tissue culture media (TCM), or 1mM AMD3100 (CXCR4i) at 4 and 24 hours are shown as an average of two replicate wells per patient per condition ±Ls.d. **g, h,** Quantitation of migrated PBMCs (**g**) or percent CXCR4^+^ cells (**h**) from patient T19 (pre-treated with 1mM AMD3100 in blue) incubated in T-cell media (TCM) with or without PDTO (30,000), and in absence or presence of 1 μg/mL CXCL12 (n=3; each) ±Ls.d at 4 hours. Statistical differences were determined using averages (for n > 1) or individual values by paired t test (a, c, d) and Student t test (f-h).

Since the CXCL12-CXCR4 chemokine axis had previously been shown to modulate T-cell migration, we sought to examine the effects of CXCR4 inhibition on PBMCs [19, 21]. Using a transwell migration assay (**Fig. 1e**) we found that pretreatment with AMD3100 significantly enhanced PBMC migration from the upper to lower chamber at 4 hours (**Fig. 1f, left**), though this effect was diminished at 24 hours (**Fig. 1f, right**). Next, we assessed whether the introduction of autologous PDTOs could impact PBMC migration at 4 hours. We confirmed that neither T-cell media nor organoid media, both of which contain growth factors/cytokines, impacted PBMC migration (**Fig. S2a**), and then proceeded with transwell migration assays. PMBC migration was augmented by pretreatment with CXCR4i, in the absence or presence of PDTOs (**Fig. 1g**). When autologous PDTOs were placed in the lower chamber of the transwell assay, PBMC migration was further increased by mean change of 4-fold (p=0.003), suggestive of a role of chemoattractants (**Fig. 1g**). We hypothesized that the amount of CXCL12 secreted from these PDTO lines was insufficient to suppress migration and that other pro-migratory factors dominated the phenotype. To study this further, we explored the effects of adding 1mM CXCL12 to the media and counterintuitively found that migration of PMBCs increased further (**Fig. 1g**). Recognizing that the PMBCs isolated from patients represent a mixed population of cells, we focused on CXCR4-expressing PBMCs and found that CXCL12 suppressed migration in this population regardless of the presence or absence of PDTOs (**Fig. 1h**). Furthermore, CXCL12-dependent suppression of migration of CXCR4-expressing lymphocytes was negated when PBMCs were pre-treated with ADM3100 in the absence of PDTOs, and to a lesser extent in the presence of PDTOs (**Fig. 1h)**.

Finally, we collected media used to culture PDTOs and tumor-adjacent pancreas PDOs for 4 days, and added this conditioned media to the transwell assay containing patient-matched PMBCs. We found that the PDTO media increased PBMC migration whereas medium from the tumor-adjacent pancreas PDO line had no effect (**Fig. S2b**). Furthermore, when compared to untreated PBMCs, AMD3100 pretreated PBMCs increased PBMC migration regardless of the conditioned medium (**Fig. S2b**). Together, these data support the role of CXCL12-CXCR4 axis in mediating immunosuppression through its effects on PBMC migration.

### Combining CXCR4 inhibition with immune checkpoint blockade and gemcitabine slows tumor progression and improves survival *in vivo*

The combination of AMD3100 and ICB has previously been shown to control tumor growth in the autochthonous *LSL-Kras^G12D/+^; LSL-Trp53^R172H/+^; Pdx-1-Cre* (KPC) PDAC mouse model [19]. Given the role of gemcitabine as a chemotherapy backbone in the current standard of care therapy for PDAC patients, and our findings that it may contribute to T-cell activation **(Fig. 1d)**, we hypothesized that combining it with AMD3100 and ICB would enhance tumor control in PDAC [27]. To investigate this, we conducted a seven-arm survival study testing gemcitabine, AMD3100 (CXCR4i), and ICB (mDX-400; mouse α-PD1 monoclonal antibody) as single agents, and as double and triple combinations in tumor-bearing KPC mice **(Fig. 2a)**. A contemporaneous cohort of vehicle-treated control KPC mice is included as reference [28]. We used three-dimensional high resolution ultrasound to monitor tumor volumes at baseline and twice weekly after starting treatment once *de novo* PDAC tumors reached 4-7 mm in diameter **(Fig. 1a)** [29]. We confirmed pharmacodynamic response to CXCR4 inhibition, by measuring white blood cell (WBC) counts at time of necropsy. As expected, compared to controls, mice receiving AMD3100 developed significant leukocytosis in keeping with AMD3100’s ability to mobilize stem cells **(Fig 2b, Fig. S3a, b)**. In contrast, mice treated with gemcitabine exhibited leukopenia, consistent with chemotherapy-induced bone marrow suppression **(Fig 2b, Fig. S3a, c)**.

**Fig. 2.**
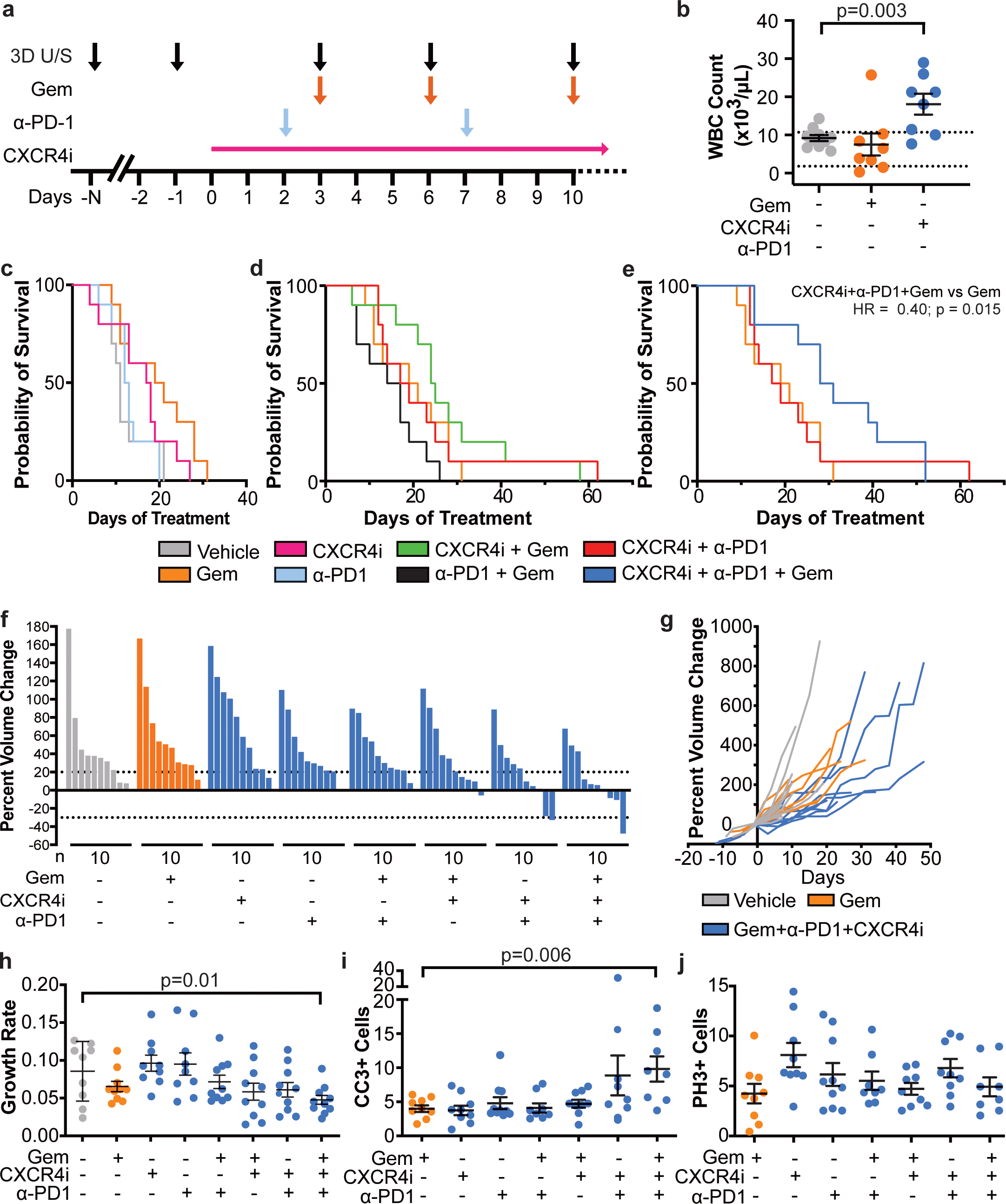
Increased survival and improved tumor control in KPC mice with the combination of gemcitabine, CXCR4 and immune checkpoint inhibition. **a**, Survival Study Schema. KPC mice were randomly assigned to one of seven cohorts (n=10 per group) and treated with various combinations of the CXCR4 inhibitor (CXCR4i) AMD3100 (via a 14-day continuous subcutaneous Azlet pump), mDX400, an αPD-1 IgG antibody, or IgG control antibody (via intraperitoneal (IP) injection) every 5 days beginning on day 2 (for a maximum of 5 doses), gemcitabine or saline control twice weekly (via IP injection) beginning on day 3. Tumor volume was measured by 3D ultrasound twice weekly until endpoint criteria were met, at which time mice were sacrificed and tumors harvested. **b**, Aggregated WBC counts for mice treated with control (grey), gemcitabine (orange), or CXCR4i therapy (blue). **c, d, e**, Kaplan-Meier survival curves depicting overall survival of mice in each treatment group, vehicle, and single agents (**c**); gemcitabine and all possible doublet combinations (**d**); gemcitabine, CXCR4i + αPD-1, and the triple combination (**e**). **f,** Best response for each tumor by mouse, based on maximal percentage of tumor reduction from baseline. **g**, Tumor growth curves depicting change in tumor volume over time for each mouse in the vehicle, gemcitabine, and triple therapy group. **h**, Tumor growth rates for each treatment cohort. **i, j**, Quantification of anti-cleaved caspase-3 (CC3) (**i**) and anti-phospho-histone 3 (PHH3) (**j**) positive tumor cells, stained using IHC, in tissue procured at time of death. Statistical differences were determined by Student’s t test (b, h, i), and log-rank test (e).

Mice on each arm of the study tolerated their respective treatments well, without unexpected early treatment-related deaths. Mice either expired due to hemorrhagic ascites, likely due to vascular invasion by the tumor, clinical failure, or cachexia **(Fig. S4)**. Mice receiving gemcitabine experienced more weight loss, likely from chemotherapy-induced anorexia **(Fig. S4b, c, f, g)**. Consistent with prior reports, gemcitabine monotherapy had a limited impact on overall survival **(Fig. 2c)** [30]. Mice treated with α-PD1 mAb monotherapy derived no significant survival benefit and those treated with CXCR4i monotherapy fared similar to gemcitabine alone **(Fig. 2c)**. Aside from a trend in improved median overall survival with CXCR4i and gemcitabine, none of the two-drug combinations showed a significant improvement in survival over gemcitabine monotherapy **(Fig. 2d)**. In contrast, the triple combination of CXCR4i/α-PD1/gemcitabine significantly increased overall survival by 48% over gemcitabine alone (p=0.0148) **(Fig. 2e)**. Notably, CXCR4i/α-PD1 therapy resulted in survival similar to mice treated with gemcitabine alone.

To better characterize tumor response to treatment, we quantified KPC tumor volumes using 3D ultrasound and compared changes between treatment groups. Relative to gemcitabine-treated mice and contemporaneous control mice, CXCR4i/α-PD1/gemcitabine-treated tumors grew more slowly, with occasional stabilizations and one frank regression **(Fig. 2f & g, Fig. S5)** [28]. Using an established algorithm to quantify tumor growth rate, we found that tumors from CXCR4i/α-PD1/gemcitabine-treated mice had a significantly lower growth rate than vehicle control treated mice (p=0.01) **(Fig. 2h)** [31]. Finally, immunohistochemistry (IHC) performed on tumor samples obtained at time of necropsy revealed increased apoptosis (cleaved caspase 3) in the CXCR4i/α-PD1/gemcitabine-treated tumors compared to gemcitabine alone (p=0.006), although no difference in mitotic rate (phospho-Histone H3) was observed.

### CXCR4i/**α**-PD1/gemcitabine combination therapy provokes immune effector cells *in vivo*

In order to capture the early effects of treatment on the KPC tumor immune microenvironment, we also performed a short-term intervention study which involved pre-treatment laparoscopic tumor biopsies paired with necropsy samples collected after 6 days of treatment **(Fig. 3a)**. Paired tumor samples from baseline biopsy and at time of necropsy (day 6) demonstrated increased apoptosis (cleaved caspase 3) (p=0.02) and decreased proliferation (phospho-Histone H3) (p=0.03) in tumors of mice that were treated with triple therapy, consistent with the reduced overall growth rate observed in the survival study (**Fig. 3b-d**). In contrast, gemcitabine monotherapy treatment only induced an increase in apoptosis (p=0.03) and had no effect on tumor growth kinetics (**Fig. S5)**, similar to our previously reported findings [30]. To investigate the effect of these distinct therapies on the tumor immune microenvironment, we interrogated the paired tumor samples for changes in natural killer (NK) and B-cells using IHC (**Fig. 3b, e&f**; NKp46 and CD19, respectively), and other immune cell subsets using quantitative multiplex immunofluorescence (qmIF; **Fig. 4**). We found that the frequency of NK cells increased in necropsy samples following treatment with CXCR4i/α-PD1/gemcitabine (p=0.03), but not in vehicle- or gemcitabine-treated controls (**Fig. 3e)**. Notably, B-cell infiltration universally increased in necropsy samples independent of treatment perhaps reflecting a response to the biopsy wound (**Fig. 3f**).

**Fig. 3.**
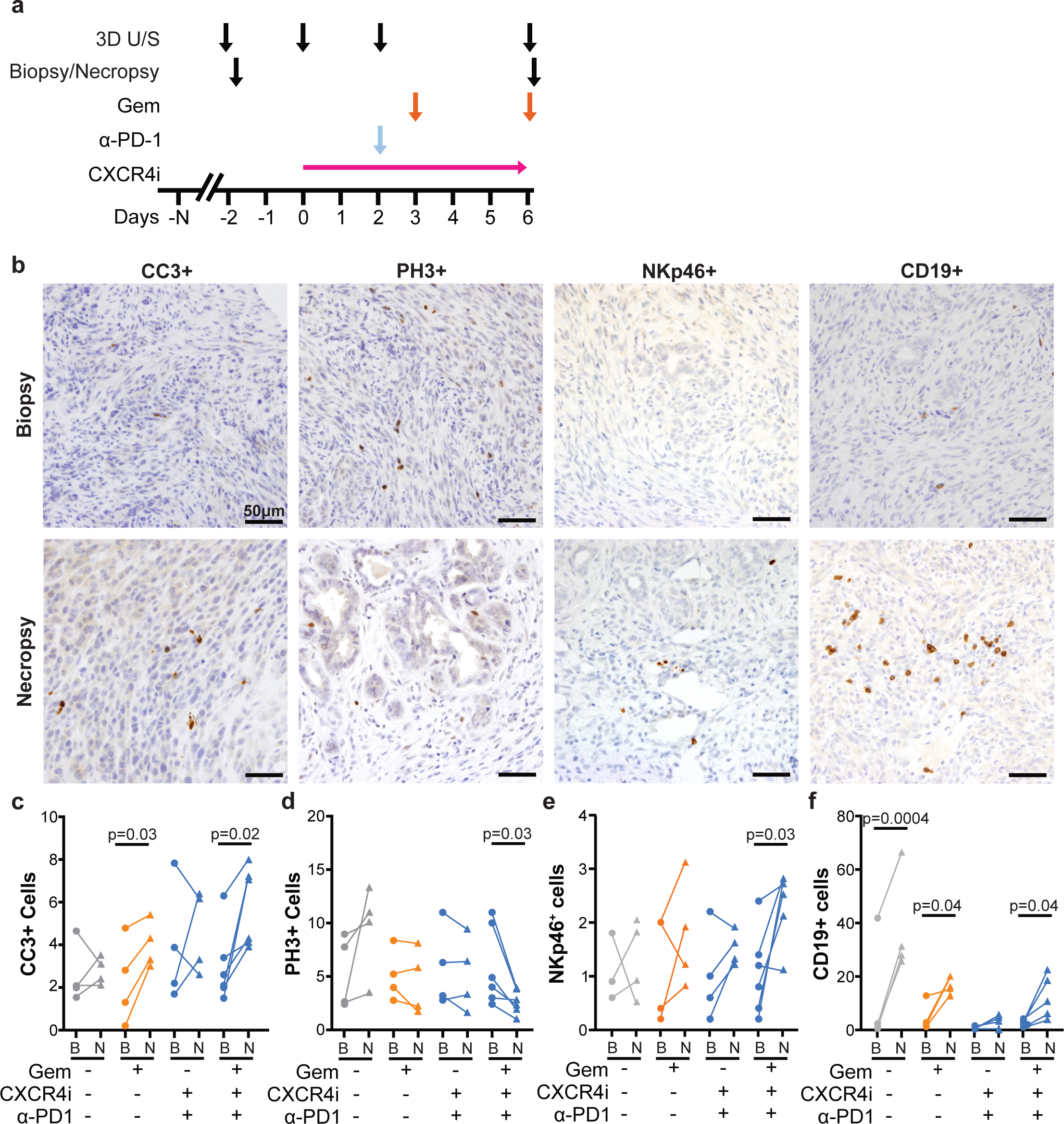
The combination of CXCR4 inhibition, immune checkpoint blockade, and gemcitabine resulted in increased apoptosis and increased natural killer (NK) and B-cell infiltration into PDAC tumors. **a**, Biopsy-Necropsy Study Schema. KPC mice were randomly assigned to one of seven cohorts (n=4 per group, except for the triple combination [n=6]) and treated with various combinations of the CXCR4 inhibitor (CXCR4i) AMD3100 (via 7-day continuous subcutaneous Azlet pump), mDX400, an αPD-1 IgG antibody, (via intraperitoneal (IP) injection) on day 2, and gemcitabine on days 3 and 6 or saline control (via IP injection). Tumor volumes were measured by 3D ultrasound. **b**, Representative images of tumor samples obtained from mice prior to starting therapy (biopsy) and at time of necropsy stained with anti-CC3, -PH3, -NKp46, and -CD19 antibodies (40x field). **c-f**, Quantitation of CC3 (**c**), PH3 (**d**), NKp46 (**e**), and CD19 (**f**) cell staining on paired pre- and on-treatment biopsy (B) and necropsy (N) samples obtained from mice treated with vehicle control, gemcitabine, CXCR4i and anti-PD1 therapy, and the triple combination. Statistical differences were determined by paired t test (c-f).

**Fig. 4.**
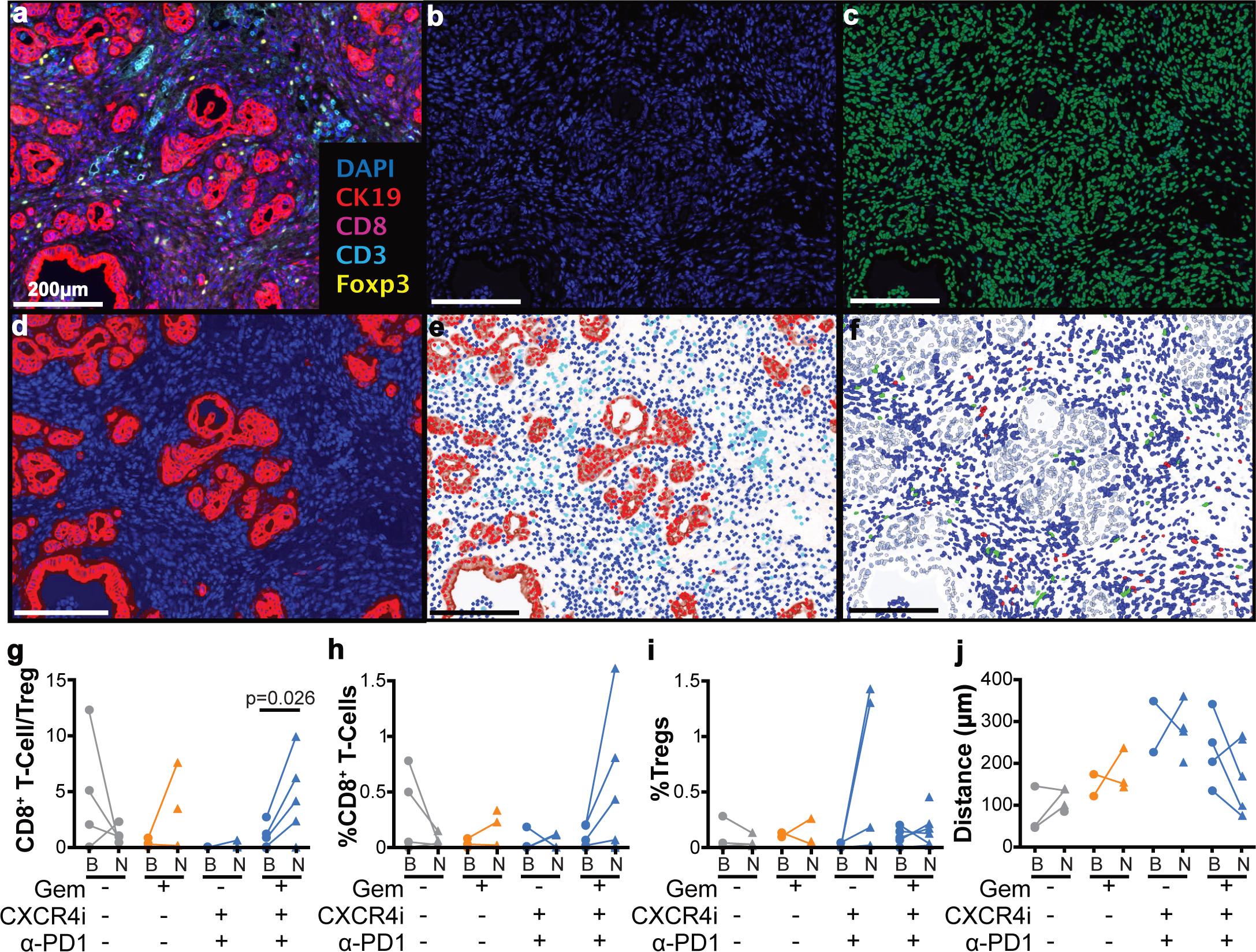
CXCR4 inhibition in combination with immune checkpoint blockade and gemcitabine resulted in an increased ratio of CD8^+^ to T-regulatory tumor-infiltrating lymphocytes. Quantitative multiplex immunofluorescence (qmIF) on biopsy and necropsy samples was performed. **a**, A representative multiplex image of necropsy tissue from a mouse treated with CXCR4i + αPD-1 + gemcitabine stained with DAPI (nuclei, blue), CK19 (tumor, red), CD3^+^ T cells (cyan), CD8^+^ T cells (magenta), and FoxP3^+^ (T-regulatory cells, yellow) is shown. **b**, DAPI staining alone. **c-f**, Representative steps in analysis using inForm® software (Akoya Biosciences®): **c**, cell segmentation; **d**, tissue segmentation; **e**, phenotyping showing base phenotypes of CD3^+^ T cells (cyan), tumor (red), and other (blue); **f**, scoring with CD8^+^ cells (red) and Foxp3^+^ cells (green). **g**, Ratio of total CD8^+^ to CD3^+^FoxP3^+^ cells in biopsy (B) and necropsy (N) specimens obtained from mice treated with various combinations of vehicle, gemcitabine, CXCR4i, and α-PD1 therapy. **h, i**, Average percentages of infiltrating CD8^+^ lymphocytes (**h**) and regulatory-T cells (CD3^+^FoxP3^+^) (**i**) per total cell population. **j**, Distance analysis using nearest-neighbor distances (assessing the closest CD8^+^ T cell to each tumor cell) showing tumor-to-CD8 distances in paired pre- and on-treatment biopsy-necropsy samples obtained from mice treated with various combination of vehicle, gemcitabine, CXCR4i, and α-PD1 therapy. Statistical differences were determined by paired t test (g).

We used qmIF to simultaneously quantify and spatially characterize tumor epithelial cells (CK19), T-cells (CD3 and CD8), and regulatory T-cells (Foxp3) in pre-treatment biopsy and on-treatment necropsy tissue samples (**Fig. 4**). Images were captured using the Vectra Multispectral Imaging System and then analyzed using the companion inForm software (**Fig. 4a-f**). We found a statistically significant increase in the CD8^+^ T-cell to Treg ratio in mice treated with triple therapy (p=0.026) (**Fig. 4g**). There was a non-significant increase in CD8^+^ T-cell infiltration in samples obtained from mice in the triple therapy cohort (**Fig. 4h**) and no statistical difference was noted in regulatory T-cell or macrophage cell density in paired tumor samples from any of the treatment groups (**Fig. 6i**, data not shown). Finally, given our observations in PDIO co-culture system on the effects of CXCR4 inhibition on T-cell migration, we used Nearest Neighbor Distance (NND) analysis to assess the clustering between cells. NND assesses the average distance between each tumor cell and its closest T-cell, and we found that T-cells trended to cluster more closely around malignant tumor cells after treatment with CXCR4i/α-PD1/gemcitabine (mean fold change of 0.676, p=0.217; **Fig. 4j**).

## DISCUSSION

CXCR4 is a widely expressed chemokine receptor on immune cells in the PDAC TME [32-34]. Furthermore, its ligand, CXCL12, is highly expressed by abundant stromal carcinoma associated fibroblasts which coat cancer cells within the PDAC TME [35, 36]. In tumors of PDAC patients, high levels of intra-tumoral CXLC12 correlates with poor prognosis [32-34, 37]. CXCL12 forms covalent heterodimers with keratin-19 on tumor cells facilitated by transglutaminase-2, and interruption of this interaction allows intra-tumoral accumulation of T-cells in mouse PDAC models [38]. Binding of CXCL12 to CXCR4 not only results in T-cell exclusion, but also in decreased T-cell activity. However, inhibition of CXCR4 when combined with ICB has been shown to reverse this process and induce tumor responses in preclinical models [19, 21, 39, 40].

To more authentically model complex autocrine and paracrine signaling between human PDAC and key effector immune cells, we developed a modified PDTO system. Unlike traditional co-culture systems that use PDAC cell lines or allogeneic PMBCs, our assay combines primary PDTOs and matched PBMCs derived from the same patient. Once established, we used this approach to interrogate key signaling pathways as well as immune cell activation and migration. We demonstrated that co-culturing PBMCs with PDTOs, but not organoids derived from tumor-adjacent tissue, increased activation of T-cells. We also noted that this activity is blunted with HLA-neutralizing antibodies and augmented with the addition of gemcitabine. These findings suggest that the increase in T-cell activation observed in the presence of PDTOs may be dependent on neo-antigen presentation.

In addition to being able to evaluate antineoplastic immunomodulators, our PDIO system can serve as a platform to dissect the role of individual targets within distinct compartments of the TME (tumor using PDTOs, stroma using CAFs, or immune using PBMCs). To assess the effects of CXLC12-CXCR4 signaling modulation on immune cell migration, we adapted our PDIO model into a transwell migration assay. We found that inhibition of CXCR4 resulted in increased PBMC migration and that introducing PDTOs into the system further augmented this activity evoking a potential role of chemoattractants released by PDTOs. Counterintuitively, CXCL12 increased PBMC migration, and this was more pronounced in the presence of PDTOs. In contrast, CXLC12 suppressed migration of CXCR4^+^ lymphocytes within this PBMC population, and this affect was rescued when PMBCs were pretreated with AMD3100. It is possible that inhibition of CXCR4 unmasked CXCR4-mediated paracrine signaling that enhanced migration of at least a subset of PBMCs. Similarly, the presence of PDTOs had distinct effects on PBMCs and CXCR4^+^ lymphocyte migration suggesting complex paracrine interactions between the tumor and the immune compartments. Although a powerful system, PDIOs are limited by scarce material (tumor tissue and PBMCs), growth and passage of PDTOs, in particular for tumor adjacent normal organoids, all of which restrict the number of conditions and replicates that can be tested.

Given our findings that treatment of PDTOs with gemcitabine enhanced CD8^+^ T-cell activation, we tested combination of CXCR4i, gemcitabine, and α-PD1 in the KPC mouse model and found that only mice treated with triple therapy experienced a survival benefit. Surprisingly, the combination of CXCR4i and α-PD1 did not prolong survival despite transiently inducing tumor stabilization as reported previously [19]. The addition of gemcitabine to CXCR4i/α-PD1 was required to decrease tumor growth rate, increase apoptotic cell death, and decrease cell proliferation. Our short-term biopsy-necropsy study demonstrated that addition of gemcitabine to CXCR4i/α-PD1 also resulted in a favorable tumor immune microenvironment, as measured by CD8^+^ T-cell/Treg ratio and a trend towards improved proximity of CD8^+^ T-cells around neoplastic cells. Combined, these results suggest that the addition of gemcitabine enhances CXCR4i/α-PD1’s ability to counter the immunosuppressive TME and increase neoplastic cell death to improve outcomes.

Our preclinical findings are consistent with those reported from clinical trials that have tested CXCR4i with or without ICB in PDAC. A study evaluating continuous 7-day infusion of AMD3100 in 26 patients and a trial testing the combination of ulocuplumab (α-CXCR4) and nivolumab (α-PD1) in 27 patients failed to demonstrate radiographic responses [40, 41]. Two additional clinical trials combined CXCR4i with ICB. None of the 14 patients treated with motixafortide with atezolizumab (NCT03193190), and only one of 29 patients treated with motixafortide with pembrolizumab experienced a radiographic partial response [22]. Finally, the COMBAT/KEYNOTE-202 trial tested the combination of 5-FU and liposomal irinotecan (5-FU/nal-I) with motixafortide/pembrolizumab and reported a modest confirmed overall response rate of 13.2% and a median PFS of 3.8 months [23].

These findings evoke critical questions regarding targeting the CXCL12-CXCR4 pathway. First, why does the combination of CXCR4i and ICB, specifically with motixafortide and pembrolizumab, appear unable to improve PFS despite eliciting an inflamed tumor immune microenvironment in PDAC patients [22]? Second, why did the addition of 5-FU and liposomal irinotecan (5-FU/nal-I) to motixafortide and pembrolizumab not result in improved outcomes [23]? We speculate that although CXCR4i/αPD1 inflames the TME, resistance develops rapidly resulting in loss of tumor control as seen in our preclinical study. One possible explanation for the limited efficacy with 5-FU/nal-I/motixafortide/pembrolizumab is that it was tested in patients who had received prior gemcitabine-containing therapy. This may have altered the TME making it less susceptible to agents targeting the CXCL12-CXCR4 axis. Another explanation is that 5-FU-based chemotherapy many not be the ideal backbone to combine with CXCR4i/αPD1. There is emerging data showing significant immunomodulatory properties with gemcitabine in PDAC. Three studies have shown that treatment with gemcitabine but not 5-FU in PDAC patients resulted in a decrease in circulating peripheral Tregs [42-44]. Furthermore, *ex vivo* IL-2-mediated induction of Tregs may be inhibited with gemcitabine, but not with 5-FU [45]. Lastly, a recent high-throughput drug screen identified gemcitabine among seven drugs that had greater toxicity for intra-tumoral Tregs than peripheral Tregs [46]. Furthermore, gemcitabine treatment inhibited tumor growth in immunocompetent but not immunocompromised allografts [46]. Gemcitabine treatment in a syngeneic subcutaneous MC38 colon adenocarcinoma model demonstrated selective depletion of tumor-infiltrating Tregs [46]. These preclinical and clinical studies demonstrate that gemcitabine has immune modulating properties and is likely better suited as the chemotherapy partner to combine with CXCR4i and ICB. Our preclinical findings provided the justification to study the combination of CXCR4i (motixafortide), αPD1 (cemiplimab), gemcitabine, and *nab*-paclitaxel in treatment-naïve metastatic pancreatic adenocarcinoma which currently is actively accruing participants (NCT04543071).

## CONCLUSIONS

The CXCL12/CXCR4 axis plays a critical role in PDAC, dampening anti-tumor immune responses and creating an immunosuppressive tumor immune microenvironment. Using a novel *ex vivo* human PDIO autologous co-culture mobilization assay and the KPC pancreatic cancer mouse model we demonstrate that the combination of CXCR4i, α-PD1 mAb, and gemcitabine therapy exerts tumor control through partial relief of immunosuppression and concordant activation of immune effector cells.

## METHODS

### Human subjects

Peripheral blood and tumor tissue were obtained from histologically confirmed treatment-naïve pancreatic ductal adenocarcinoma patients. Patients provided written informed consent to Research Ethics Committee-approved protocol (Institutional Review Board at the United States Center), in compliance with Good Clinical Practice, local regulatory requirements, and legal requirements (CUIMC IRB AAAR9431).

### Organoid culture

Tumor tissue obtained from surgical resection from PDAC patients was processed within 24 hours, cut into small pieces and enzymatically digested using 1.5 mg/mL collagenase II (Sigma-Aldrich), 10 μg/mL hyaluronidase type IV (Sigma-Aldrich) and 10 μM Y-27632 (Sigma-Aldrich) before embedding cells in Geltrex. After Geltrex solidification for 20 minutes at 37 °C and 5% CO_2_, cells were overlaid with human pancreatic cancer organoid medium. Organoid medium is composed of Ad-DF+++ (Advanced DMEM/F12 (Gibco) supplemented with 2mM Ultraglutamine I (Lonza), 10 mM HEPES (Gibco), and 100U/ml Pencillin/Streptomycin (Gibco)), 10% Noggin-conditioned medium, 20% R-spondin1-conditioned medium, 1x B27 supplement without vitamin A (Gibco), 1.25mM N-Acetylcysteine (Sigma-Aldrich), 10mM nicotinamide (Sigma-Aldrich), 50 ng/mL human recombinant EGF (Peprotech), 500nM A83-01 (Tocris), 3μM SB202190 (Cayman Chemicals) and 10nM prostaglandin E2 (Cayman Chemicals). In the first two weeks of organoid culture, 1x Primocin (Invivogen) was added to prevent microbial contamination. Organoids were passaged approximately every two weeks by mechanical dissociation and washed with PBS to dissociate organoids to single cells and replate in fresh Geltrex. Organoids passages 10-20 were used in the experiments.

### Isolation of Peripheral Blood Mononuclear Cells (PBMCs) and plasma by Ficoll-Paque density gradient centrifugation

ACD-A-anticoagulated blood was centrifuged at 2000rpm for 10 min and the top layer containing plasma was removed and aliquoted into 6 vials for cryopreservation at -80°C. The remaining blood was diluted with phosphate-buffered saline, pH 7.4 (PBS) and layered over 15 ml of the Ficoll-Paque PLUS (Fisher Scientific, Cat. No.: 45-001-749). Gradients were centrifuged at 1800 rpm for 30 min at room temperature in a swinging-bucket rotor without the brake applied. The PBMC interface was carefully removed by pipetting and washed with PBS by centrifugation at 2000rpm for 10 min twice. PBMCs were cryopreserved in fetal bovine serum (FBS; Fisher Scientific, Cat. No.: 10-099-14) containing 10 % dimethyl sulfoxide (DMSO; Fisher Scientific, Cat. No.: 14-190-144) and stored in liquid nitrogen until required for downstream analyses.

### Patient-derived organoids/autologous PBMC co-culture assay

This assay was conducted according to M. Cattaneo’s protocol [47]. Culture media for PBMC was composed of RPMI 1640 (Gibco), supplemented with 2mM Ultraglutamine I, 1:100 penicillin/streptomycin and 10% male human AB serum (Sigma-Aldrich) (“T-cell medium”). One day before co-culture, autologous PBMCs were thawed in pre-warmed (37

°C) T cell medium (human serum was replaced with FCS during thawing) and incubated for 15 minutes with 25U/mL benzonase (Merck). After washing, cells were resuspended at 2-3X10^6^ cells/mL in T-cell medium supplemented with 150U/mL IL-2 and cultured overnight at 37°C. Prior to co-culture, tumor organoids were isolated from Geltrex (the basement membrane matrix) to avoid the presence of non-human protein material in the co-cultures, dissociated to single cells with TrypLE Express, resuspended in T-cell medium and pre-stimulated overnight with 200ng/mL human recombinant IFNγ (Peprotech) to maximize antigen presentation. 96-well U-bottom plates were coated with 5μg/mL anti-CD28 (clone CD28.2, eBioscience) and kept overnight at 4°C. The next day, anti-CD28-coated plates were washed twice with PBS and PBMCs were seeded at a density of 10^5^ cells/well and stimulated with single cell-dissociated organoids at a 20:1 effector:target ratio. Co-cultures were performed in the presence of 20 μg/mL anti-PD1 (Nivolumab, Merus/Selleckchem) to counteract the induction of PD-L1 by IFNγ, 150 U/mL IL2 (Proleukin, Novartis) to support T-cell proliferation and plate-bound anti-CD28 antibodies (CD28.2; eBioscience) to provide co-stimulation. Half of the medium, including IL-2 and anti-PD-1, was refreshed two to three times per week. Every week, PBMCs were collected, counted and replated at 10^5^ cells/well, and re-stimulated with fresh tumor organoids. T-cell reactivity was assessed after 14 days of co-culture by flow cytometry. Briefly, 10^5^ PBMCs were restimulated with tumor organoids at a 2:1 effector:target ratio and seeded in anti-CD28-coated plates in the presence of 20 μg/mL anti-PD-1 and co-cultured for 5 hours. Mouse anti-human CD107 α-PE antibodies (BD) were added at the start of co-culture. Golgi-Plug (1:1000, BD) and Golgi-Stop (1:1500, BD) was added after 1 hour and co-culture continued for an additional 4 hours. Cells were washed twice in FACS buffer and stained with the following antibodies: anti-CD3-PerCP-Cy5.5 (BD), anti-CD4-FITC (BD), anti-CD8-BV421 (BD), and near-IR viability dye (Life technologies) for 30 minutes at 4 °C. Cells were washed twice in FACS buffer, fixed and stained for intracellular IFNγ (anti-IFNγ-APC, BD) using the Cytofix/Cytoperm kit (BD), according to manufacturer’s instructions. PBMC stimulated with 50 ng/mL phorbol 12-myristate 13-acetate (PMA, Sigma-Aldrich) and 1 μg/mL ionomycin (Sigma-Aldrich) served as positive controls and PBMC cultured without tumor stimulation as negative controls. For HLA-neutralizing co-culture assay, dissociated PDTOs were incubated with 150 μg/mL of mouse anti-human HLA-ABC (clone W6/32, ThermoFisher) to block MHC-I and 30 μg/mL mouse anti-human HLA-DR/DP/DQ (clone Tu39, BD Biosciences) to block MHC-II for 30 minutes at 37 °C. PDTOs were treated with 10 μM gemcitabine for 16 hours prior to co-culture at 37 °C.

### Transwell migration assay

To assess the chemoattractant properties of tumor organoids on patient’s PBMC, *ex vivo* cell transwell migration assays were established. Human cryopreserved PBMCs from patient with PDAC were thawed, washed twice and resuspended in T-cell migration media (RPMI + 1% FBS) with or without recombinant human CXCL12 (R&D Systems, 350-NS). PBMC (50,000) suspension in 100μL was loaded in the upper wells of Boyden chamber plates (Corning Transwell Supplier no. 3422; 8-μm membrane pore size). Lower chambers contained 600μL T-cell migration media with or without 30,000 single-cell patient-derived organoids and in the presence or absence of 1μg/mL of recombinant human CXCL12. To evaluate the effect of CXCR4 inhibition on migration, PBMCs were pretreated with 1mM AMD3100 (Sigma, A5602) on ice for 30 min then washed twice with PBS and loaded in the upper chambers of Boyden chamber plates. Migration was allowed for the allocated time at 37 °C and 5% CO_2_, Transwell inserts were then removed and cells that had migrated into the lower chamber were counted using a hematocytometer.

### Flow Cytometry

Human PBMCs migrated to the bottom wells of Boyden chamber plates, were collected, counted, washed in PBS, blocked with FACS buffer (PBS 1% FBS), and incubated for one hour at 4°C with fluorescent antibodies, according to manufacture protocols: CD3 (Per-CP-Cy5.5-conjugated, eBioscience 332771), CD8 (BV421-conjugated, BD 562429), CD4 (FITC-conjugated, BD 555346), CXCR4 (AF700 and PE-conjugated, respectively clones 12G5 and 1D9, BD Pharmingen), CD11b (FITC-conjugated, BD 562793), CD56 (PerCpCy5.5-conjugated, BD 560842), FoxP3 (PE-conjugated, BD 560046. After washing twice, cells were fixed in PFA 2% for 20 min, washed, and acquired on a Fortessa BD LSRII Cell Analyzer (Becton Dickinson).

### Mice and Imaging Procedures

All murine studies were approved by the Columbia University institutional animal care and use committee (IACUC) and conducted in accordance with the NIH “Guide for the Care and Use of Laboratory Animals.” *Kras*^LSL.G12D/+^; *P53*^LSL.R172H/+^; *Pdx1-Cre* (KPC) mice were generated as has been previously described [48]. KPC mice were monitored for tumor formation by weekly palpation. Upon positive tumor palpation, tumors were monitored with twice weekly ultrasound until an average cross-sectional diameter of 4-7 mm was reached. Upon reaching this size, mice with tumors in the head of the pancreas were enrolled randomly into one of the seven treatment arms in the survival study. Those mice with tumors in the tail of the pancreas were randomized into one of the seven treatment arms in the biopsy study. Tumor growth was monitored twice weekly after enrollment and volume quantification was carried out as previously described [29].

### Survival study

The survival study consisted of seven treatment arms with ten mice per arm. Each arm consisted of either single agent or any of all possible combinations of the CXCR4 inhibitor (AMD3100, subcutaneous Alzet pump administered continuously from day 0 at 48.7 mg/ml), α-PD1 (mDX400 IgG from Merck Inc, intraperitoneal (i.p.) injection starting on day 2 and given every 5 days to a max of 5 doses at 10 mg/kg) and gemcitabine (i.p. starting day 3, and given twice weekly at 100 mg/kg) (**Fig. 2a**). Mice were euthanized once they reached endpoint criteria consisting of a combined physical and behavioral metric designed in consultation with Columbia IACUC. Overall survival was defined as the time from randomization until endpoint criteria were met.

### Biopsy study

For the biopsy study, the same treatment combinations were used but enrolled four mice per arm, except the triple therapy arm which had six mice. Subsequently, four additional KPC mice were enrolled on a vehicle arm, as controls. Mice were euthanized six days after initiating treatment. Biopsy surgeries were carried out as previously described in accordance to Columbia University IACUC standards and protocol [49]. Blood samples were taken by cardiac puncture at time of necropsy.

### Immunohistochemistry

Tissue for immunohistochemistry was fixed at room temperature overnight in 10% phosphate-buffered formalin, transferred to 70% ethanol, and then processed, prior to being embedded in paraffin. Tissues were sectioned to 5 µm thickness and baked onto positively charged slides at 60°C for 30 minutes. Slides were deparaffinized with xylene and hydrated through graded concentrations of alcohol. Sections were then subjected to heat-induced epitope retrieval with 10mM citrate buffer for 5 minutes using a pressure cooker. Slides were then cooled to room temperature ∼15 minutes in ice bath. Endogenous peroxidases were quenched by immersion in 3% hydrogen peroxide for 20 minutes. Slides were blocked for 1 hour at room temperature in 1.5% normal horse serum (Vector Laboratories, S-200) + 2% Animal Free Blocker (Vector Laboratories, SP-5030) in 0.1% PBS – Tween20. Slides were then stained with the appropriate primary antibody diluted in blocking solution and incubated overnight at 4°C: Phospho-Histone H3 (Cell Signaling, #9701, 1:200), Cleaved Caspase 3 (Cell Signaling, #9664S, 1:1000), CD19 (Abcam ab245235, 1:800), NKp46 (R&D, #AF2225, 1:800). The following day, slides were incubated for 30 minutes with the appropriate secondary antibody: ImmPRESS HRP Anti-Rabbit IgG secondary (Vector Laboratories, MP-7401). Immunodetection was performed using the HRP system and visualized with chromogen 3,3’-diaminobenzidine (Vector Laboratories, SK-4105). Slides were then dehydrated and counterstained with hematoxylin.

Images of each stained slide were obtained using a Leica LMD6500 microscope. Quantification for CC3 and PHH3 was done in a blinded fashion, counting positive cells per 40x field for 10 fields per sample. For CD19 and Nkp46, the 10 most dense areas with tumor on it at 40x magnification field sections were chosen blindly by an independent pathologist using QuPath v0.2.2. Cells shed into the lumen or close to the biopsy scar were excluded from evaluation. Quantification was then done manually and expressed as the mean for each slide.

### Quantitative Immunofluorescent Staining

Full-section 5-mm slides of tissue specimens were stained using Opal multiplex 6-plex kits, according to the manufacturer’s protocol (PerkinElmer®, now Akoya Biosciences®), for DAPI, CD3 (clone SP7; Spring Bioscience; 1:100 dilution), CD8 (clone 4SM15; eBioscience; 1:2000); CK19 (clone EPNCIR127B; Abcam; 1:450), FoxP3 (clone FJK-16s; eBioscience; 1:100 dilution), and F4/80 (clone SP115; Spring Bioscience; 1:100); PD-L1 (clone 22C3; Dako; 1:150). Briefly, the opal multiplexing method involves serial immunofluorescent labeling of specific epitopes through the use of covalently bound tyramide signal amplification [50, 51]. During this process, epitope-specific primary and secondary antibody complexes are added and then subsequently removed, while leaving behind a covalently-bound fluorescent signal. Controls included single stains and an unstained slide for each batch of stained slides.

### Multispectral imaging

Prior to staining, H&E slides were viewed by a gastrointestinal pathologist to determine representative areas for multispectral image capture at 20 magnification using VECTRA® (PerkinElmer, now Akoya Biosciences). As capturing lymph node or lymphoid aggregates would potentially skew results, the gastrointestinal pathologist was tasked with identifying up to 10 representative areas per slide predominantly composed of tumor. These images were factored equally into the analysis for each mouse. Using a multispectral camera, VECTRA® is able to capture spectral information from a multiplexed panel of targets. These images then undergo spectral unmixing, a process which requires each fluorophore to be identified by using representative single-stained slides for each antibody and a representative autofluorescence spectrum from an unstained sample. From this process, images from each single-stained and unstained slides were used to create a multispectral library in inForm®. Finally, intensity of each fluorescent target was extracted from the multispectral data using linear unmixing.

### Image analysis

All images were analyzed using inForm® software. DAPI counterstaining was used to differentiate cellular and nuclear compartments, with each associated cytosolic or membrane bound protein detected via presence of a specific stain (FoxP3, CD3, CD8, F4/80, PD-L1, and CK19). Tissue segmentation was performed by highlighting examples of CK19+ tumor and non-tumor tissue, and through an iterative process, the computer algorithm was able to “learn” each tissue type and segment based on the presence of the CK19 marker and morphology (**Fig. 4d**). Following this, cell segmentation was performed using the minimum DAPI signal to accurately locate all cells and adjusted based on splitting, in order to avoid hypersegmentation or hyposegmentation of each nucleus, and adjusting sizes of nuclei to fit both tumor and immune cells (**Fig. 4c**). Cells were phenotyped by using the phenotyping step of inForm® software. Approximately 10 representative cells for each base variable were chosen to train the phenotyping algorithm: tumor (CK19, red dots), T cells (CD3, cyan dots), and other (negative for CK19, CD3, PD-L1, and F4/80, blue dots) (**Fig. 4e**). Finally, the images were scored for intensity based on each individual base variable and the secondary marker for further phenotyping of CD8, FOXP3 and (**Fig. 4f**). Data obtained from all representative images was compiled to yield values for each mouse. Image data were exported from inForm version 2.2.1 (PerkinElmer) and processed in separate software designed in RStudio (version 0.99.896; https://github.com/thmshrt/transform_essential). In this software, images were combined and analyzed to calculate numbers of each cell type.

### Spatial clustering analysis

To assess clustering in immunofluorescence imaging nearest neighbor distances (NNDs) were computed using the spatstat R package [52]. NNDs were calculated by finding the closest CD8^+^ T-cell to each tumor cell, and taking the median of this distance across all tumor cells in the sample.

## Acknowledgements

KPO received support for this work from the NIH National Cancer Institute (3 R01 CA157980-04) as well as through a sponsored research agreement from Merck. GAM received the Conquer Cancer Foundation Young Investigator Award (2015) and the Herbert Irving Comprehensive Cancer Center Velocity Award (2019).

## Supplemental figures

**Fig. S1. Expression of CXCL12 and CXCR4 in PDTOs and PBMCs. a**, Quantification of CXCL12 secreted by PDTOs once confluency was achieved. Data are presented as an average of technical replicates (n=3) ±Ls.d for three PDTO lines (hT56, T18, T19) generated from individual PDAC patients, and are assessed by Mesoscale Discovery. **b**, CXCR4 surface expression on PBMCs before and after 30 minutes of treatment with AMD3100, using αCXCR4 (clone 12G5) and DAPI counterstaining, as observed by immunofluorescence. **c**, CXCR4 and CXCL12 expression in single-cell RNA sequencing data from in peripheral blood immune cells (left panel) and cells found in pancreatic cancer tissue (right panel) [53]. For each cell type in each tissue source, circle size represents the fraction of the respective cell type relative to all cells profiled. Circle color indicates scaled average expression where scaling considers both tissue sources. Statistical differences were determined by Student’s t test (a).

**Fig. S2. Neither T-cell media nor organoid media impact PBMC migration. a,** Quantification of PBMC migration, following pre-treatment with either 1mM AMD3100 (grey bars) or PBS (white bars), to the bottom chambers of Boyden transwell assay containing either T-cell media (TCM), organoid growth media (OGM), or a combination of TCM and OGM 1:1 (TCM/OGM). **b**, Quantification of PBMC migration, following pre-treatment with either 1mM AMD3100 (grey bars) or PBS (white bars), to the bottom chambers of Boyden transwell assay containing PDTO or adjacent normal tissue PDTO supernatant. Number of PBMCs which migrated to the lower chamber was measured after 4 hours. Each condition was performed in duplicate wells and reported as averages.

**Fig. S3. Aggregated white blood cell (WBC) counts for all treatment groups in the survival study. a,** Quantification of WBC counts for each treatment group. **b**, WBC counts for mice receiving (+) or not receiving (-) a CXCR4i-containing treatment regimen (open data points indicate mice that also received gemcitabine). **c**, WBC counts for mice receiving (+) or not receiving (-) a gemcitabine-containing treatment regimen (open data points indicate mice that also received CXCR4i therapy). Statistical differences were determined by Student’s t test (c).

**Fig. S4. Survival Weights and Cause of Death for all treatment groups in the survival study.** Charts depicting the proximal cause of death assigned for KPC mice in each of treatment groups, calculated as described in Methods. Clinical failure was used to describe a constellation of symptoms including malignant (clear) ascites, jaundice, loss of temperature regulation, and inactivity. The change in body mass for KPC mice over time is depicted.

**Fig. S5. Tumor volume change over time for single agents and doublets compared with gemcitabine.** Graph showing tumor volumes of mice treated with gemcitabine (orange) versus CXCR4i, α-PD1, and all possible doublet combinations are plotted versus the mouse’s age in days.

