## Supplemental Figures for "Cytotoxic chemotherapy potentiates the immune response and efficacy of combination CXCR4/PD-1 inhibition in models of pancreatic ductal adenocarcinoma"

### Slide 1
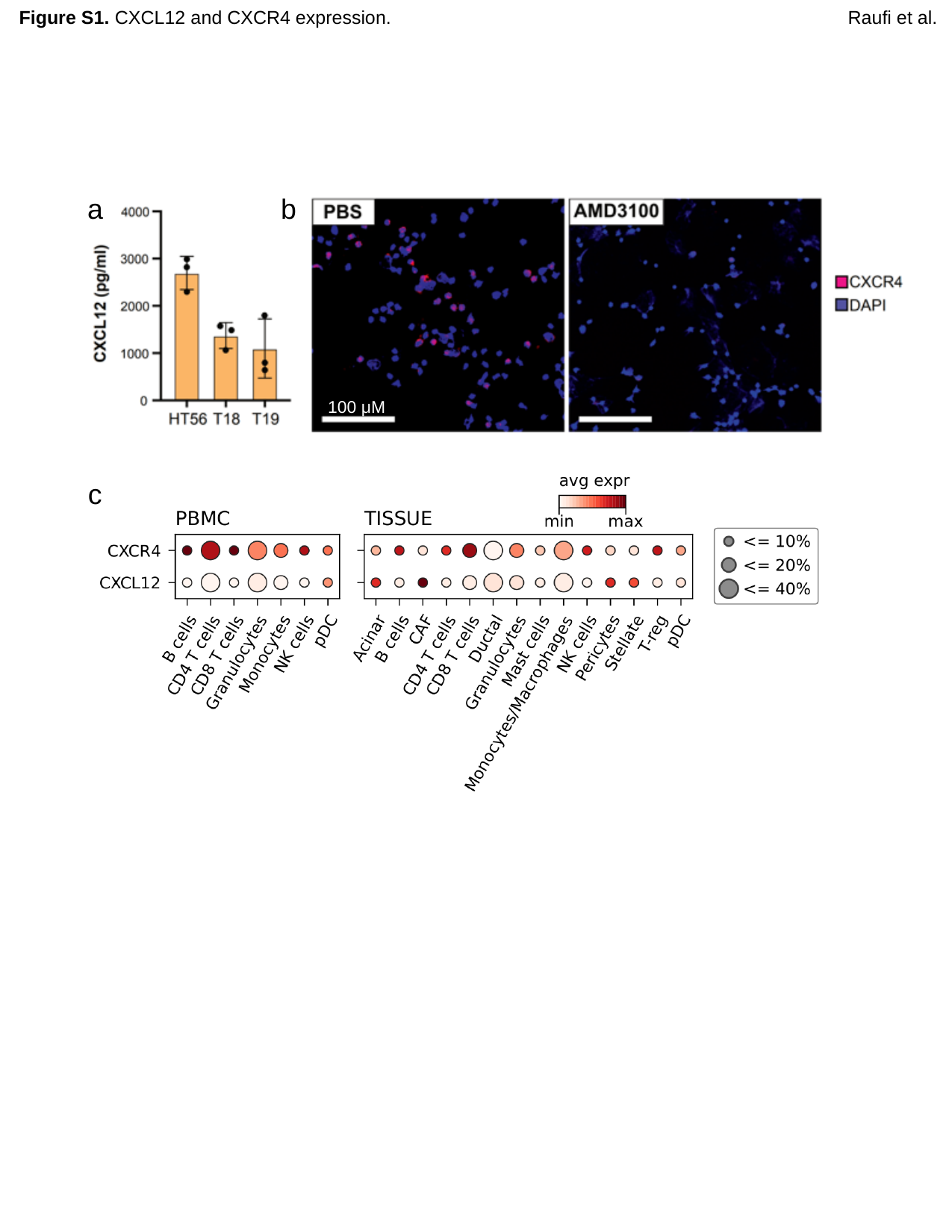

Figure S1. CXCL12 and CXCR4 expression.
Raufi et al.
a
b
100 μM
c
c

### Slide 2
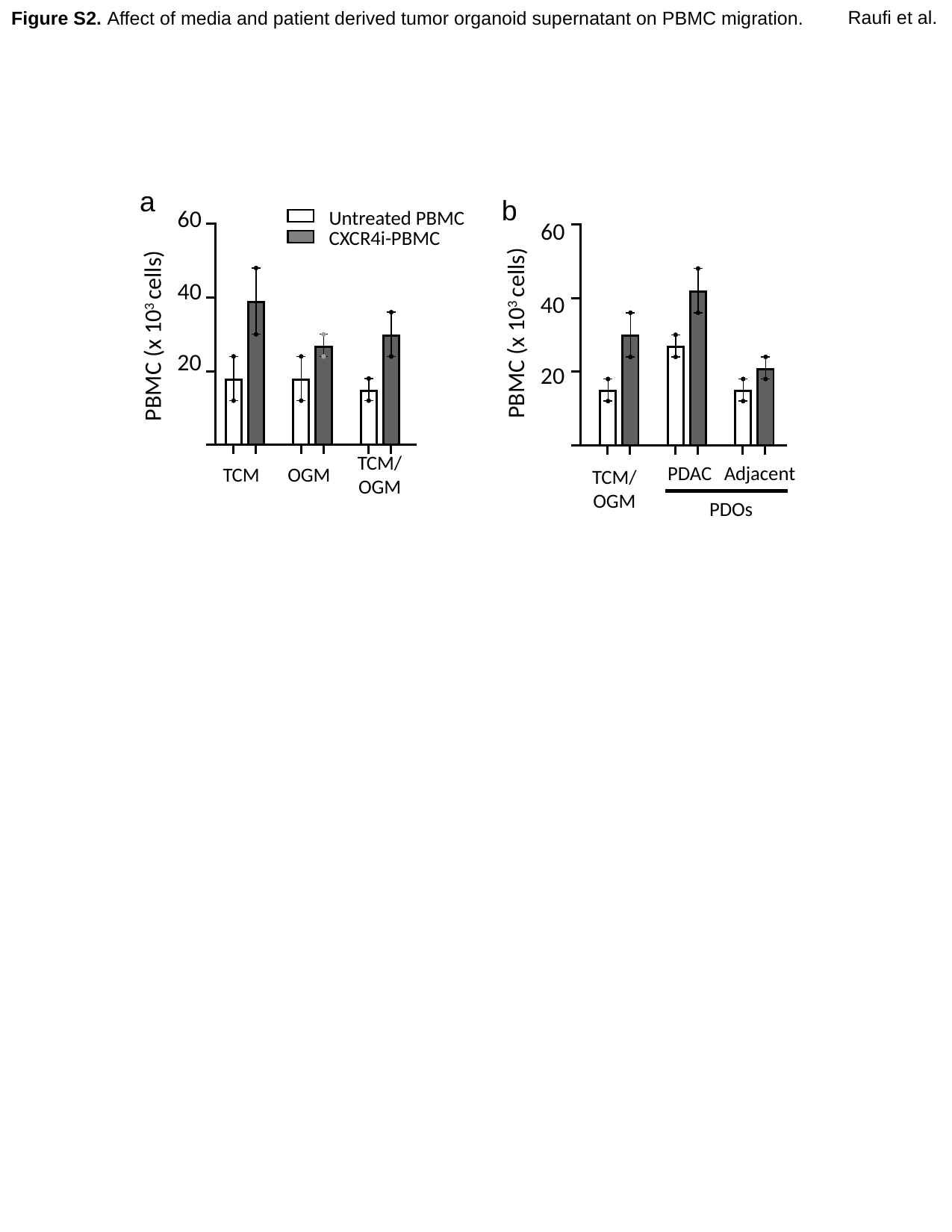

Raufi et al.
Figure S2. Affect of media and patient derived tumor organoid supernatant on PBMC migration.
a
b
60
 Untreated PBMC
60
 CXCR4i-PBMC
40
40
PBMC (x 10^3^ cells)
PBMC (x 10^3^ cells)
20
20
TCM/
OGM
PDAC
Adjacent
TCM
OGM
TCM/
OGM
PDOs

### Slide 3
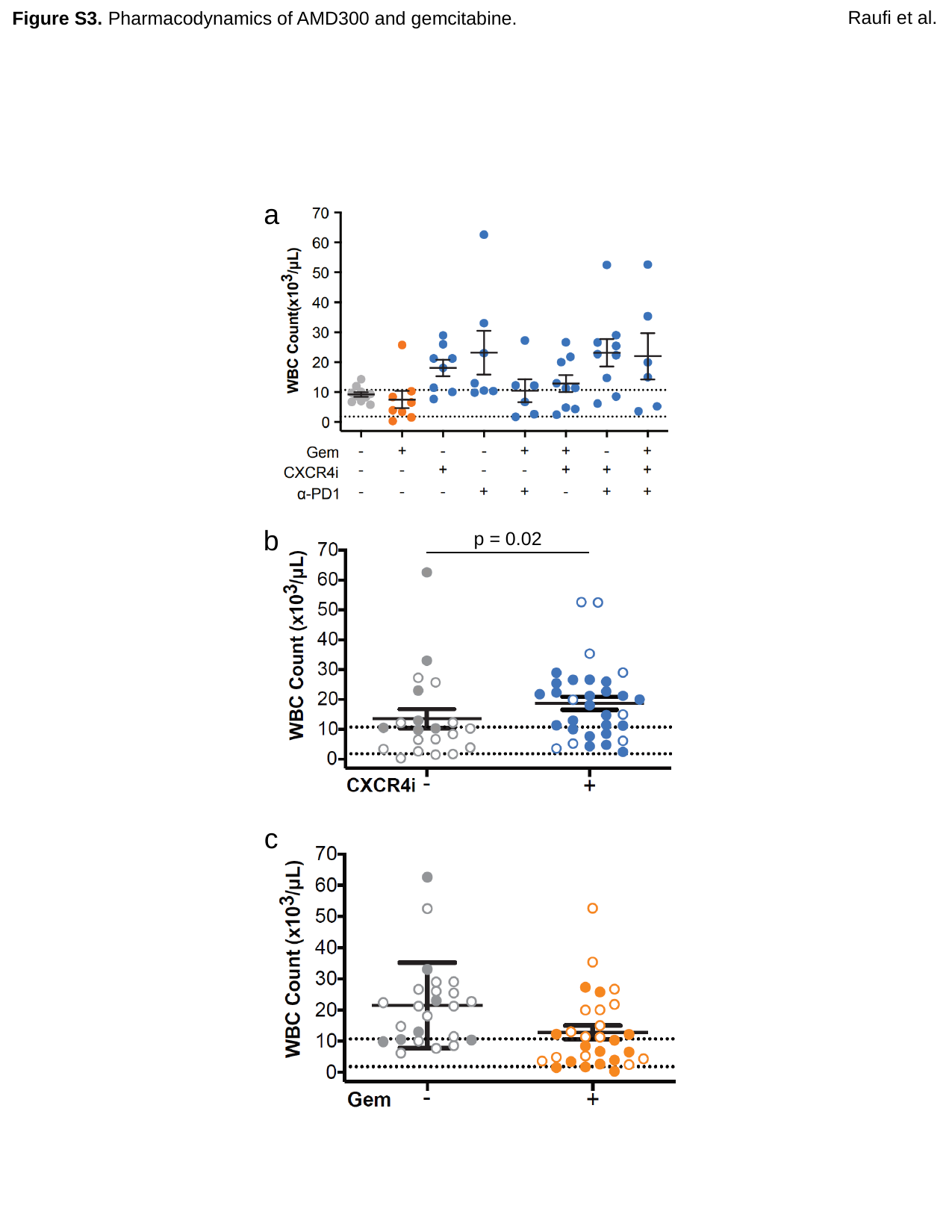

Raufi et al.
Figure S3. Pharmacodynamics of AMD300 and gemcitabine.
a
b
p = 0.02
c

### Slide 4
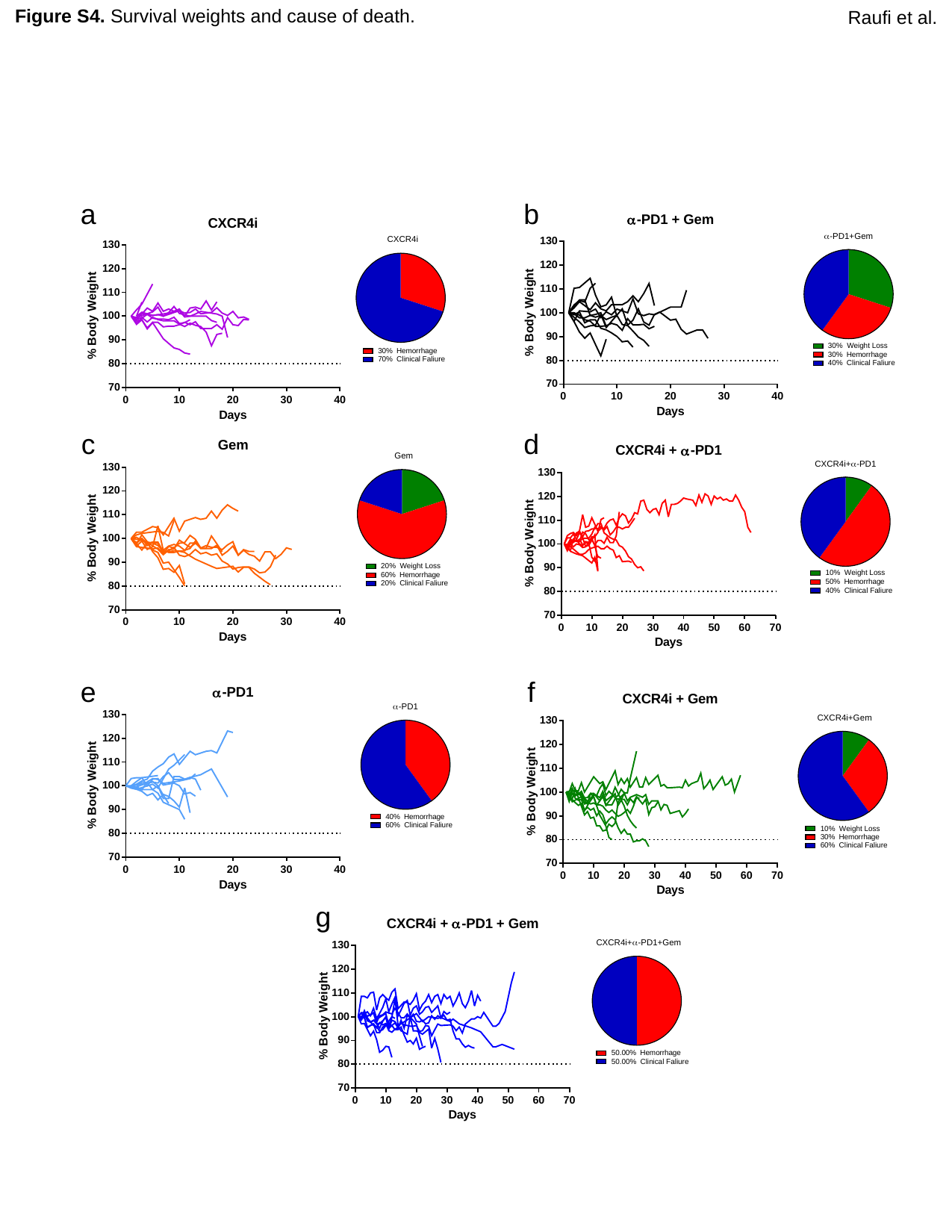

Raufi et al.
Figure S4. Survival weights and cause of death.
a
b
c
d
e
f
g

### Slide 5
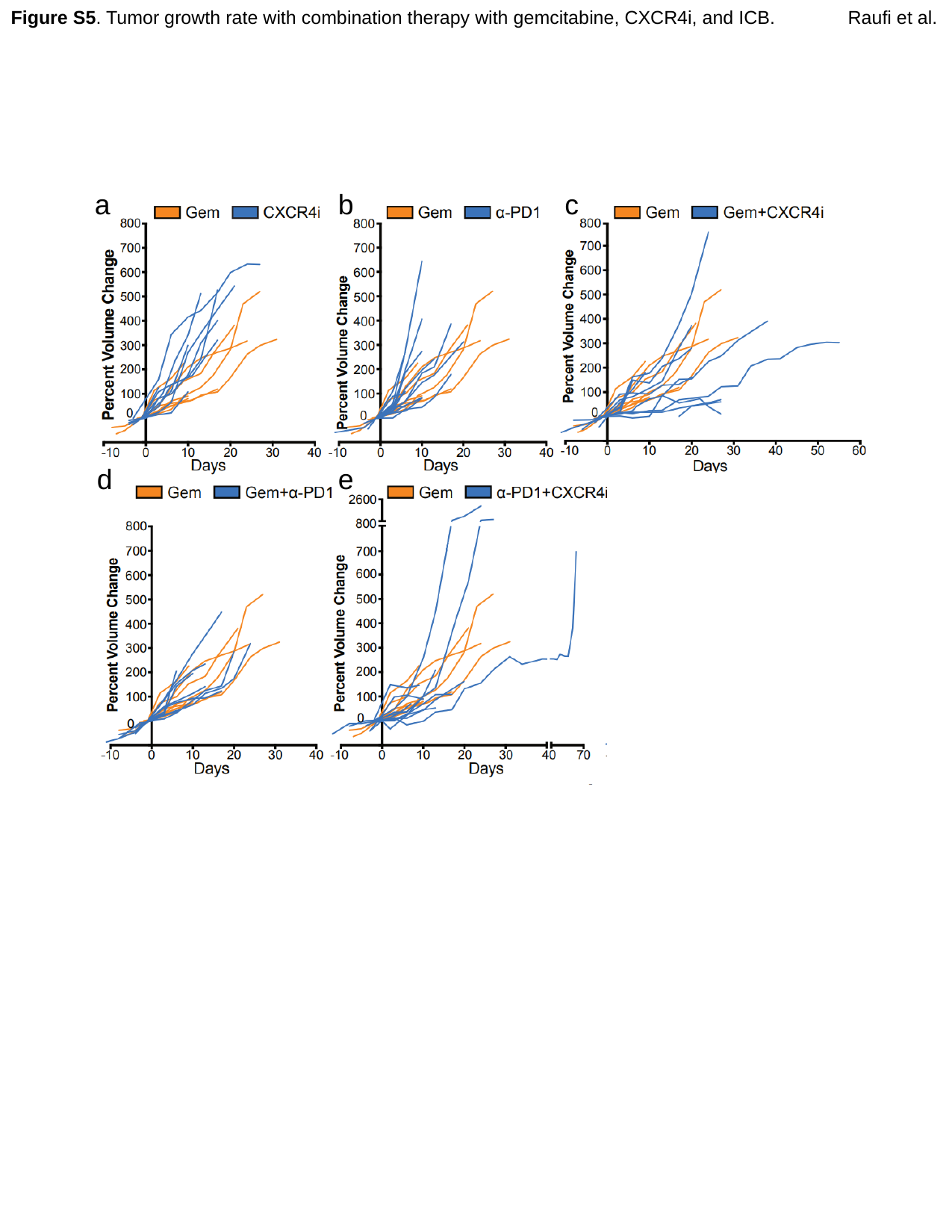

Figure S5. Tumor growth rate with combination therapy with gemcitabine, CXCR4i, and ICB.
Raufi et al.
b
c
a
d
e
